# Multiscale Computational Modeling of the Cardiopulmonary Consequences of Postnatal Hyperoxia with Implications for Preterm Born Children

**DOI:** 10.1101/2025.10.01.679825

**Authors:** Salla M Kim, Filip Jezek, Pim JA Oomen, Gregory P Barton, Feng Gu, Daniel A Beard, Kara N Goss, Mitchel J Colebank, Naomi C Chesler

## Abstract

Moderate to extreme preterm birth (<32 weeks gestation) affects cardiopulmonary structure and function and is associated with increased risk of heart failure through adulthood. The rat hyperoxia (Hx) model (term born; postnatal Hx exposure) captures biventricular changes, including at the cell- and organ-scale, and pulmonary vascular remodeling seen in preterm humans. However, synthesizing these measures across scales and organ systems is challenging. We hypothesized that in-silico modeling of biventricular mitochondrial, myofiber, and organ-scale function plus circulatory function could capture key features of cardiopulmonary abnormalities due to preterm birth. Therefore, we calibrated a multiscale model to subject-specific biventricular pressure-volume data previously obtained from Hx rats alongside normoxic (Nx) controls to investigate the abnormalities in cardiopulmonary function at multiple scales in this animal model of human preterm birth. The calibrated model demonstrates excellent agreement with the data and captures the expected pulmonary vascular changes and right ventricular dilation seen in preterm born children. Our multiscale modeling approach captures cardiopulmonary abnormalities across spatial scales and provides an innovative approach to explore the consequences of preterm birth beyond preclinical experimental data alone. This is a foundational step in understanding the impact of preterm birth on cardiopulmonary disease in childhood as well as adulthood.

## 1. Introduction

Preterm birth (less than 37 weeks gestation) affects 1 in 10 live births in the United States (Bensley et al., 2010; Ferré et al., 2016). Neonatal care has improved over the last several decades, leading to an increased number of moderately to extremely preterm (less than 32 weeks gestation) born babies surviving into adulthood, with the consequence that we are only now beginning to understand the short- and long-term cardiopulmonary effects. The combination of the preterm newborn’s underdeveloped cardiopulmonary system and their frequent need for life-saving respiratory support such as oxygen supplementation places them at high risk for cardiopulmonary injury (Bavineni et al., 2019). The cardiopulmonary consequences of preterm birth continue past infancy (Perez et al., 2020); through adulthood, those born premature are at increased risk for systemic and pulmonary hypertension, with an up to 17-fold increased risk for heart failure, cardiometabolic diseases, and stroke (Carr et al., 2017).

Clinically, the effects of preterm birth on cardiopulmonary structure and function span multiple spatial scales. At the organ and tissue scales these effects include reduced biventricular chamber sizes and microvascular density, ventricular hypertrophy and fibrosis, and changes to contractile function (Barton et al., 2021; François et al., 2022; Sixtus et al., 2024). Additionally, preterm neonates are exposed to a relatively hyperoxic (Hx) environment, and since mitochondrial DNA is very susceptible to oxidative damage, they are at high risk for cardiopulmonary mitochondrial dysfunction (Ali et al., 2024; Bavineni et al., 2019; Mohammadi et al., 2022). Coupled with high right ventricular (RV) afterload caused by an under-developed pulmonary circulation, these multiscale changes contribute to impaired RV-pulmonary vascular coupling (Dartora et al., 2021; Greer et al., 2021; Mulchrone et al., 2020).

Animal models can provide valuable functional data at multiple scales. The lung developmental stage at term for rodents is comparable to moderately to extremely preterm humans, and exposing rat pups born at term to 10-14 days of postnatal Hx (∼85%) has been shown to capture key cardiopulmonary features seen in human preterm birth and is among the most used rat models of preterm birth (Berger & Bhandari, 2014; DeFreitas et al., 2024; O’Reilly & Thébaud, 2014; Ravizzoni Dartora et al., 2022; Velten et al., 2010). At postnatal day 21 (P21), which corresponds to early childhood, rat pups born at term exposed to 14 days of postnatal Hx have been shown to suffer from pulmonary hypertension, RV dysfunction, depressed cardiac mitochondrial oxidative capacity, and RV-pulmonary vascular uncoupling (Kumari 2019). These findings correspond to clinical findings in preterm-born children and provide useful insight into the disease state (Arjaans et al., 2022; Jeong et al., 2024; Kumari et al., 2021; Mulchrone et al., 2020). However, disparate measurements across multiple organ systems and scales can be challenging to synthesize. Mechanistic computational modeling can provide insights into the underlying mechanisms of the observed cardiopulmonary dysfunction and simulate physiological outputs that are beyond the available data, encouraging the generation of new hypotheses and experimental designs. Furthermore, subject-specific modeling can provide insight into disease progression in a single individual and generate an enhanced set of biomarkers for contrasting control and disease groups or identifying new disease subgroups (Colunga et al., 2020; Gu et al., 2025; Jones et al., 2021; Kachabi et al., 2025). Combining experimental data with multiscale modeling can extend our understanding of complex processes, but also reduce the need for numerous, costly experiments that may be infeasible.

In this study, we tested our hypothesis that in-silico, subject-specific modeling of biventricular mitochondrial, myofiber, and organ-scale function and circulatory function could capture key features of the cardiopulmonary abnormalities due to preterm birth. With multiscale data previously collected in the postnatal Hx rodent model of preterm birth (Kumari et al., 2019; Patel et al., 2017), we simulated cardiopulmonary function using a multiscale cardiac model (Marzban et al., 2020) that includes the biventricular “TriSeg” heart model (Lumens et al., 2009) with calcium-dependent active myofiber force, and a closed-loop lumped-parameter circulation model as presented in Kim et al. (2023). We developed a workflow for subject-specific, multiscale modeling and analysis of pediatric, Hx rodents, including subject-specific sensitivity analyses and calibration of influential parameters to biventricular pressure-volume data. Overall, we confirmed our hypothesis that the model can represent RV and pulmonary vascular adaptations that are consistent with experimental and clinical findings. Our research provides a framework that combines literature and available experimental data to identify key mechanisms of cardiopulmonary dysfunction across spatial scales due to postnatal hyperoxia in rats. The model codes are publicly available to facilitate further independent studies.

## 2. Methods

### 2.1 Animal Data

Data collected from 12 rats (5F, 7M) exposed to neonatal hyperoxia as a model of preterm birth and 7 control rats (2F, 5M; normoxia only) are used here (Kumari et al., 2019). As detailed in Kumari et al. (2019), timed Sprague-Dawley dams delivered naturally to term. Within 12 hours of birth, pups were randomly exposed to either normal room air (Nx; 21% oxygen) or oxygen rich air (Hx; 85% oxygen) for 14 days. The Hx conditions mimic the relatively hyperoxic environment experienced by humans born preterm as previously demonstrated (Berger & Bhandari, 2014; Velten et al., 2010). The dams were rotated between chambers during the exposure period to avoid maternal oxygen toxicity. At P21, hemodynamic data including left ventricular (LV) and RV pressures and volumes were measured using invasive catheterizations and analyzed on Notocord Systems software (Notocord Systems, Croissy Sur Seine, France). The catheterizations were conducted via serial ventricular punctures using a 27g needle that was previously clotted (to prevent leakage) for both ventricles: the LV then the RV. Due to positioning there was no appreciable blood loss, but as an assurance a small piece of blotting paper was used to apply pressure while removing the needle and inserting the catheter.

Sequential heart beats (20-50) were selected and processed in MATLAB software. The average pressure and volume signals were used for analysis and model calibration as further described in **Section 2.5**. The data displayed in **Table 1** were used for nominal parameter estimations, the same data separated by sex are shown in **Table S1**, where other than RV end systolic volume (ESV), which was greater in the Hx females, there was no significant difference between the females and males

**Table 1.** Group Averages of Measured Data.

| Metric | Units | Nx (n = 7) | Hx (n = 12) | p |
| --- | --- | --- | --- | --- |
| Female/Male | - | 5(F)/7(M) | 2(F)/5(M) | - |
| BW | g | **49.9 ± 1.2** | **53.4 ± 2.8** | **\*8.9e-3** |
| T | s | **0.190 ± 0.044** | **0.179 ± 0.015** | **NS** |
| SV | $\mu\text{L}$ | **28 ± 9** | **28 ± 7** | **NS** |
| LV ESV | $\mu\text{L}$ | **16 ± 9** | **18 ± 8** | **NS** |
| RV ESV | $\mu\text{L}$ | **24 ± 11** | **23 ± 9** | **NS** |
| LV EDP | mmHg | **4.4 ± 2.0** | **5.1 ± 1.4** | **NS** |
| LV ESP | mmHg | **65 ± 9** | **72 ± 11** | **NS** |
| RV EDP | mmHg | **2.9 ± 1.5** | **3.8 ± 1.5** | **NS** |
| RV ESP | mmHg | **26 ± 3** | **38 ± 6** | **\*2.3e-4** |

BW – body weight; T – time; SV – stroke volume; ESV – end systolic volume; EDV – end diastolic volume; EDP – end diastolic pressure; ESP – end diastolic pressure; LV – left ventricle; RV – right ventricle.
\* Indicates a significant difference from a two-sample unpaired student's t-test ( $p < 0.05$ ). NS - not significantly different.

### 2.2 Data Processing

Because LV and RV pressure and volume readings were recorded sequentially rather than simultaneously, the measured cardiac cycle duration (T, in seconds) differed from the LV and RV due to heart rate variability. To correct this, we averaged the LV and RV cycle durations and used this single cycle duration for both ventricles. Differences between LV and RV stroke volume (SV, in µL) due to the asynchronous catheter recording were accounted for by setting a common SV for both ventricles based on the average LV stroke volume for each animal since the conductance catheter measurement is typically more accurate for the LV geometry (Wearing et al., 2025). There was less variability in the measured end-systolic volume (ESV) compared to the measured end-diastolic volume (EDV), so we used the measured RV ESV and let RV EDV = RV ESV + SV. Also, for several animal models, the recorded end-diastolic pressure was negative, a calibration error that was corrected by shifting the pressure curve upward to match the average end diastolic pressure of the remaining animals in the same condition (Nx, Hx) and sex (M, F) group. Finally, the catheter artifacts during isovolumetric contraction can occur due to catheter positioning (Wearing et al., 2025). Given that these artifacts are not physiological and do not correspond with the disease phenotype, we enforced isovolumetric contraction and relaxation in the data.

### 2.3 Multiscale Model Overview

We present an adapted multiscale cardiovascular model that includes crossbridge- and calcium (Ca^2+^)-dependent myofiber mechanics embedded within the TriSeg model and a lumped parameter circulatory model as summarized in **Figure 1**. In total there are 54 parameters and 46 differential equations, including 33 for crossbridge mechanics, 3 for myofiber mechanics, 4 for the TriSeg, and 6 for circulation dynamics. The model codes are available at https://github.com/sallakim/Multiscale-Hx-P21-Model-2025.

**Figure 1.**
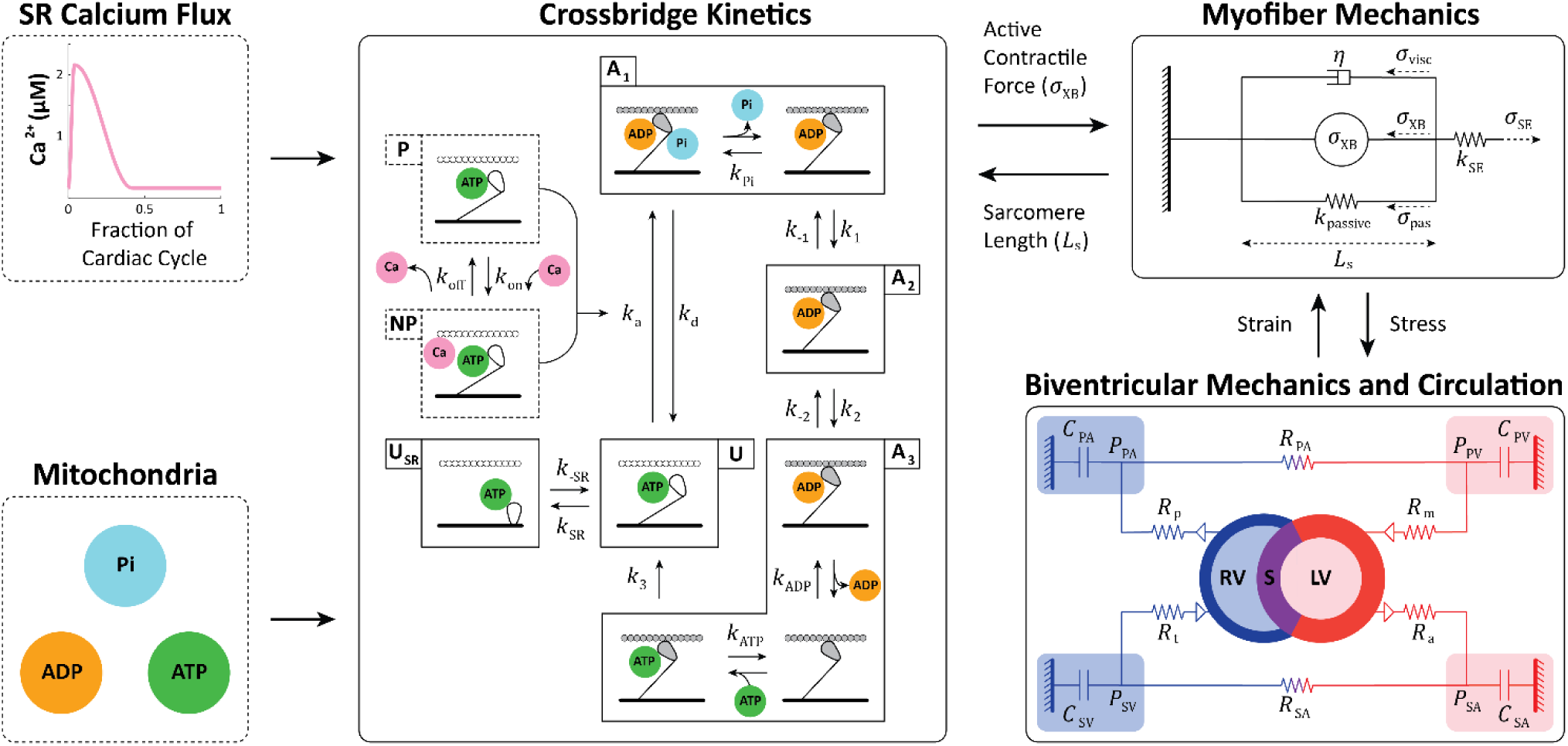
Multiscale, Multiorgan Cardiopulmonary Model Schematic. The model includes four sub-models: crossbridge kinetics, myofiber mechanics, biventricular mechanics, and circulation; with model inputs: sarcoplasmic reticulum (SR) calcium flux and mitochondria. The mitochondrial metabolites adenosine triphosphate (ATP), adenosine diphosphate (ADP), and inorganic phosphate (Pi) to determine rate constants in the crossbridge cycle, and a calcium (Ca) flux for Ca^2+^-dependent activation and active stress generation. **Crossbridge Cycling.** The five-state crossbridge sub-model has three actin and myosin attached states and two unattached states. The three attached states are 1) loosely-attached, A1, 2) strongly-attached, A2, and 3) post-ratcheted and strongly-attached, A3. The remaining states are 4) unattached, U, and 5) super-relaxed and unattached, USR. Together, these states describe crossbridge kinetics where A1 + A2 + A3 + U + USR = 1. The binding and unbinding of ATP, ADP, and Pi to myosin and the transitions between states are dictated by rate constants, *k*. The unbinding rate of Pi within state A1 is *k*_Pi_, the forward and reverse transition rate constants from state A1 to A2 are *k*_1_ and *k*_−1_. A3 state includes 3 “intermediate states” where ADP is released and ATP binds, and the rate of transitions for these intermediate states are *k*_ADP_ and *k*_ATP_, correspondingly. The forward and reverse rate constants between the unbound, relaxed (U) and the super-relaxed (USR) states are given by *k*_SR_ and *k*_−SR_, respectively. The transition between permissible, P, and non-permissible, NP, pseudo-states is mediated by the binding, *k*_on_, and unbinding, *k*_off_, of Ca^2+^ to troponin C. This transition dictates the attachment rate, *k*_a_, of myosin to actin, with a detachment rate, *k*_a_. The crossbridge kinetics determine the active contractile force, *σ*_XB_, used in the myofiber model. **Myofiber Mechanics.** The myofiber mechanics model is represented as a spring with active, *σ*_XB_, a sarcomere length-dependent passive, *σ*_pas_, viscous (*η*), *σ*_visc_, and series elastic, *σ*_SE_, force. **Biventricular Mechanics and Circulation**. The left and right ventricles (LV, RV) are represented as semi-spherical, separated by a septum (S), and comprised of thick walls according to the TriSeg model. The lumped-parameter circulation follows an electrical circuit analogy and includes the systemic arterial and venous compartments (SA, SV) and the pulmonary arterial and venous compartments (PA, PV) in addition to the LV and RV for a total of six compartments. The atrial (a), pulmonary (p), mitral (m), and tricuspid (t) valves are represented as diodes that only allow forward flow with resistance, *R*. Vascular compartments are described as compliant chambers with compliance *C*.

#### 2.3.1 Crossbridge Kinetics and Myofiber Mechanics Sub-Models

The myofiber mechanics sub-model integrates a Ca^2+^-dependent crossbridge kinetics sub-model to simulate active and passive myocardial wall stress, which has been described in detail in Marzban et al. (2020). Here, we provide an overview of the model and only detail significant changes and model components particularly relevant to this study.

A moment-based distribution approach gives a system of five ordinary differential equations that represents three attached and two unattached crossbridge states (Graham, 2020; Tewari et al., 2016). The crossbridge mechanics model links the mechanical and biochemical kinetic processes by incorporating metabolite concentrations, adenosine triphosphate [ATP], adenosine diphosphate [ADP], and inorganic phosphate [Pi] into calculations of crossbridge transition rates.

The crossbridge model includes actin and myosin model components, and the states and transitions of the crossbridge kinetics sub-model are illustrated in **Figure 1**. The three actin and myosin attached states A_1-3_ correspond to loosely attached, strongly attached, and post-ratcheted states, respectively. The remaining states are an unattached state, U, and an unattached, super-relaxed state, U_SR_. The transitions between these states are dictated by the rate constants, *k* (s^−1^). Together, these states and transition rates describe the crossbridge kinetics where A_1_ + A_2_ + A_3_ + U + U_SR_ = 1. The transition from no Ca^2+^ bound, permissible, P, and Ca^2+^ bound, non-permissible, NP, is driven by Ca^2+^ binding, *k*_on_, and unbinding, *k*_off_, to troponin C. Finally, actin-myosin attachment is described by *k*_a_, which is dependent on the Ca^2+^ dynamics.

The transition from non-permissible to permissible states is mediated by a Ca^2+^ activation function (Campbell et al., 2018). Here, we represent cytosolic Ca^2+^ concentrations as a function of time similar to other models (Chung et al., 2016; Hunter et al., 1998; Kim et al., 2023; Tran et al., 2017) with timing parameters that allow for subject-specific variability in contractile behavior, where

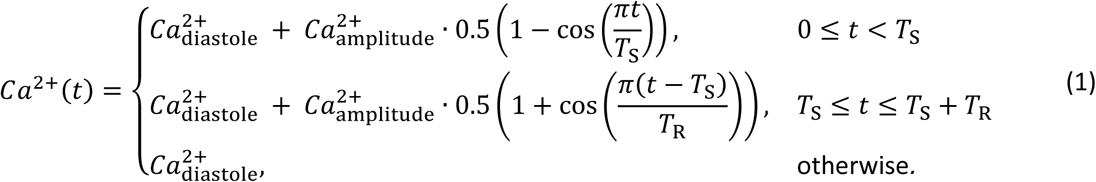

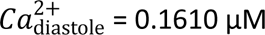 is the minimum Ca^2+^ concentration at diastole and 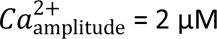 prescribes the amplitude of the sinusoidal relationship and therefore maximal Ca^2+^ concentration during systole. These values are set to correspond with the magnitudes in the data reported by (Marzban et al., 2020). The timing parameters include *T*_S_ = *k*_TS_ ⋅ *T* and *T*_R_ = *k*_TR_ ⋅ *T*, where *T* (s) is the period of one cardiac cycle and *k*_TS_ and *k*_TR_ (unitless) are the proportion of the cardiac cycle spent during increasing and decreasing Ca^2+^, respectively. These cardiac timing parameters were calibrated for each animal according to the pressure and volume time course data (see **Section 2.6**).

Ca^2+^-dependent active stress, *σ*_XB,*i*_ (kPa) where *i* = LV, RV, SEP, in the model is determined from the proportion of crossbridges in the attached, post-ratcheted state (A_3_), defined by the 0^th^ moment of state A_3_, *P*^0^ (unitless), and contributions from the attached bound, pre-ratcheted state (A_2_), according to

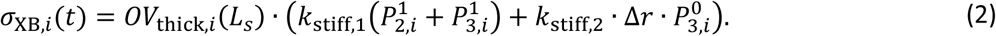

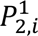 and 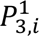 (µm) are the 1st moment state of the A_2_ and A_3_ strain probability distributions, *k*_stiff,2_ (kPa µm^−1^) is the stiffness constant due to the working stroke of the crossbridge, *k*_stiff,1_ (kPa µm^−1^) is the stiffness constant due to the myosin-actin interaction, Δ*r* (µm) is the crossbridge displacement associated with ratcheting deformation, and *OV*_thick,*i*_(unitless) is the fraction of thick filament overlap as a function of time-dependent sarcomere length, *L*_s,*i*_(µm). Additional details regarding crossbridge states can be found in Marzban et al. (2020) and Tewari et al. (2016).

We simplified the passive force, *σ*_pas,*i*_ (kPa), formulation as in Kim et al. (2023) to be represented by a single exponential relation for collagen recruitment that depends on the time-dependent sarcomere length, *L*_s,*i*_, and the contractile element length at zero active stress, *L*_sc0_ (µm). This is given by

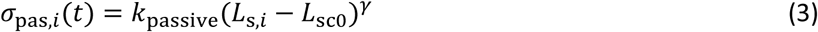

where *k*_passive_ (kPa µm^−1^) is the passive stiffness constant, and 𝛾 (dimensionless) is the steepness of the length-tension relationship.

The full myofiber mechanics model includes contributions from the active force generated by the crossbridge kinetics, passive and viscous forces, and a series element force. The overall force balance in the model is

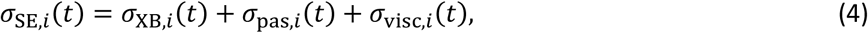

where *σ*_SE,*i*_ (kPa) is the series elastic element and *σ*_visc,*i*_ (kPa) is the viscous stress contributed from the dashpot (*η*) and determined by the rate of change of sarcomere length,

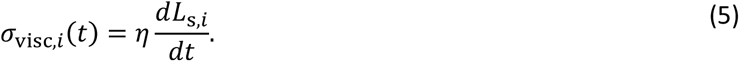

#### 2.3.2 Biventricular Mechanics and Circulation Sub-Models

For organ-scale dynamics, we model the heart and circulation as in Kim et al. (2023) using the biventricular TriSeg model (Lumens et al., 2009) and a zero-dimensional (0-D) lumped parameter circulatory model following an electrical circuit analogy with six compartments: LV, RV, systemic arteries (SA), systemic veins (SV), pulmonary arteries (PA), and pulmonary veins (PV). The pericardium is typically removed or ruptured when PV loops are obtained experimentally in rodents as in the data used here. Thus, we remove the pericardial constraint, as in Kim et al. (2023), because it is an additional complexity that does not reflect the experimental data. Briefly, the ventricles are modeled as three semi-spherical thick-walls for the LV and RV free walls and septum (SEP). Active and passive cardiac myofiber mechanics and the TriSeg geometry determine the biventricular pressures which drive flow in the circulation. The SA, SV, PA, and PV systems are represented using Windkessel models with linear compliance, and flow between the compartments is dictated by Ohm’s law with linear resistance. Finally, the mitral, aortic, tricuspid, and pulmonary valves are represented as pressure-sensitive resistances that only allow forward flow.

### 2.4 Nominal Model Parameterization for Pediatric, Hyperoxic Rats

The TriSeg and crossbridge models have been previously parameterized for adult humans, rats, and mice to study various cardiovascular disease cases (Colebank et al., 2023; Jones & Oomen, 2025; Kim et al., 2023; Lumens et al., 2009; Marzban et al., 2020; Pewowaruk et al., 2018; Tewari et al., 2016), but this is the first time these models have been parameterized for pediatric and hyperoxic rats. Crossbridge parameters from Marzban et al. (2020) were modified for early life Nx and Hx Sprague Dawley rats as described below.

#### 2.4.1 Mitochondrial Inputs

Kumari et al. (2019) reported suppressed mitochondria complex 1 activity, increased mitochondrial density, and increased oxygen consumption in the RV myocardium of Hx animals at P21. Complex 1 is crucial for ATP production, as cytosolic ATP decreases with reduced complex 1 activity. In return, ADP and Pi accumulate due to the absence of ATP. To reflect these changes in the model, we assume a 5% decrease in ATP and a 5% increase in ADP and PI concentration in Hx animals.

#### 2.4.2 Crossbridge Kinetics and Myofiber Mechanics Sub-Models Parameterization

Prakash et al. (1999) observed increased Ca^2+^ sensitivity of force with lower stiffness per area in the myocardium of Sprague Dawley neonates (P0-P3) compared to adults (P84) from skinned muscle experiments. Their findings suggest that this lower force in the neonatal myocardium is primarily from a 40% lower rate of crossbridge attachment; however, this effect may be exaggerated due to their two-state assumption. Additionally, the animals in our study are older, at P21 as compared to P0-P3. Therefore, we assume an intermediate reduction of 20% in the actin-myosin attachment rate constant, *k*_a_, for P21 to reflect the older age.

Patel et al. (2017) found that passive stiffness of isolated RV trabeculae from Hx rats at P21 doubled in comparison to the Nx controls. Correspondingly, we increased the RV passive stiffness in the model, *σ*_pas,RV_, via the passive stiffness constant, *k*_passive,RV_, by a factor of two for the Hx animals.

Patel et al. (2017) also found an approximately 70% increase in the maximum Ca^2+^ activated force, 𝐹_max_, at pCa = 4.5 (negative decadic logarithm of concentration, [Ca^2+^] = 10^−4.5^ = 32 µM) in the Hx animals compared to the Nx controls, indicating an increased Ca^2+^ sensitivity of force. We conducted a model-based investigation to determine which crossbridge parameters increase 𝑝*C*𝑎_50_ and 𝐹_max_. To do this, we first simulated the force-Ca^2+^ experiment using the isolated crossbridge and myofiber models. We fixed the sarcomere length at 2.2 µm, varied Ca^2+^ concentration from pCa 4.5 to pCa 7.0 and recorded the resulting force for several parameter changes. A total of 11 parameters can increase 𝐹_max_ or 𝑝*C*𝑎_50_ to reproduce the experimentally observed changes to the force-Ca^2+^ (Patel et al., 2017) as summarized in **Table 2**. However, *k*_1_, 𝐾_D_, 𝐾_T_, [ADP], and [ATP] must be changed by orders of magnitude outside of physiological domains to achieve the experimentally observed change in 𝑝*C*𝑎_50_ and 𝐹_max_. Therefore, *k*_stiff,2_, Δ*r*, *k*_on_, *k*_off_, *k*_a_, *k*_3_, or some combination of these parameters are more likely to generate the observed changes in the myofiber contractile behavior. Further experimental data is required to determine which of these changes are responsible for the experimentally observed changes to the force-Ca^2+^ curve. In the absence of these data, we opted to decrease *k*_off_ and increase *k*_stiff,1_ and *k*_stiff,2_ to match the 1.04% leftward shift in 𝑝*C*𝑎_50_ and 1.67% increase in 𝐹_max_ with Hx (Patel et al., 2017). The changes made to the Nx crossbridge parameters to represent Hx are summarized in **Table 3**.

**Table 2.** Parameters that Reproduce Experimentally Observed Changes the Force-Calcium Curve.

| Parameters | Experimental Observation<br>(Patel et al., 2017) |
| --- | --- |
| $\uparrow k_{stiff,2}$, $\uparrow \Delta r$, or $\uparrow \uparrow k_1$ | $\uparrow F_{max}$ |
| $\uparrow k_{on}$, $\downarrow k_{off}$, or $\uparrow k_a$ | $\uparrow pCa_{50}$ |
| $\downarrow k_3$, $\downarrow \downarrow K_D$, $\uparrow \uparrow K_T$, $\downarrow \downarrow [ATP]$, or $\uparrow \uparrow [ADP]$ | $\uparrow F_{max}$ and $\uparrow pCa_{50}$ |

Model parameters that when increased ( ↑ ) or decreased ( ↓ ) yield changes to the force-calcium curve that mimic experimental observations. The number of arrows dictates the relative magnitude of changes. $k_{stiff,2}$ - stiffness constant due to working stroke of the crossbridge; $\Delta r$ - transition rate constant from loosely to strongly bound; $k_{on}$ - binding rate of $Ca^{2+}$ to Troponin C; $k_{off}$ - unbinding rate of $Ca^{2+}$ to Troponin C; $k_a$ - myosin-actin rate of attachment; $k_3$ - myosin-actin detachment rate; $K_D$ - ADP dissociation constant; $K_T$ - ATP dissociation constant; [ATP] - ATP concentration; [ADP] - ADP concentration.

**Table 3.** Hyperoxia-Induced Crossbridge Parameter Changes from Normoxia to Match Data from (Patel et al., 2017)

| Symbol | Definition | Percent Change |
| --- | --- | --- |
| [ATP] | Cytosolic concentration of ATP | -5% |
| [ADP] | Cytosolic concentration of ADP | +5% |
| [Pi] | Cytosolic concentration of Pi | -5% |
| $k_{passive,RV}$ | RV myofiber passive stiffness constant | +100% |
| $k_{off,RV}$ | RV rate constant of Ca detaching from Troponin C | -40% |
| $k_{stiff,1,RV}$ | RV stiffness constant due to myosin–actin interaction | +70% |
| $k_{stiff,2,RV}$ | RV stiffness constant due to working stroke of crossbridges | +70% |

#### 2.4.3 Biventricular Mechanics and Circulation Sub-Models Parameterization

Total blood volume is estimated by bodyweight (**Table 1**) using 0.06 mL/g of bodyweight as a reference for rodents (Lee & Blaufox, 1985). Nominal parameterizations for ventricular wall volumes and reference areas are calculated based on measured data from **Table 1**. First, to approximate the ventricular wall volumes, 𝑉_w,*i*_(cm^3^), we used measured ventricular wall weights from a previous study and assumed a cardiac wall density of 1.055 g/mL (Gheorghe et al., 2019; Vinnakota & Bassingthwaighte, 2004). Individual LV and RV wall weights were not available from the rats with hemodynamic data available used here. A cohort of P21 Nx and Hx rats that did not undergo hemodynamic measurements (Kumari et al., 2019) had their wall weights, 𝑊_*i*_, and bodyweight collected. Therefore, we calculated the average wall weight to bodyweight ratios, 𝑊*R*_*i*_, from this cohort for the LV and septum, 𝑊*R*_LV+SEP_ (Nx, n =14, 3.8 ± 0.43; Hx, n = 12, 3.2 ± 0.37), where the LV is 2/3 and the SEP is 1/3 of this wall weight, and for the RV, 𝑊*R*_RV_(Nx, n =14, 0.86 ± 0.13; Hx, n = 12, 1.00 ± 0.10). Then, given the wall weight to bodyweight ratio, we used bodyweight data to approximate the ventricular wall weights and subsequently the ventricular wall volumes for the subjects used in this present study. We approximated the midwall reference areas, 𝐴_m,ref,*i*_, based on the measured LV and RV EDV data and an assumption of spherical geometry as detailed in Kim et al. (2023). Therefore, the nominal parameter calculations for the TriSeg geometry are consistent across all subjects and the nominal parameters vary due to variations in the animal data (**Table 1**). Nominal TriSeg geometry parameters are used as initial starts for calibration as detailed in **Section 2.6**.

### 2.5 Sensitivity Analysis and Parameter Subset Selection

To determine which parameters to estimate given the available data, we conducted a sensitivity analysis to understand how variations in parameters affect the model output (Olsen et al., 2019). Key predictive outcomes of the model are LV and RV pressure and volume signals, and we are interested in identifying which parameters, relative to the available data, are most influential. Thus, we restricted our analysis to determining the influence of organ-scale and cardiac timing parameters on dynamic pressure and volume signals for both the LV and RV, which yields 18 parameters:

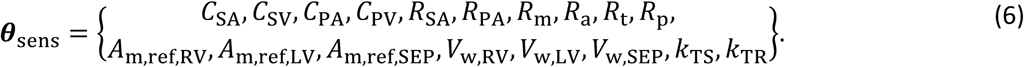

We performed a local, derivative-based sensitivity analysis to determine how perturbations, ℎ, to the parameters, 𝜽_sens_, influence the model output, 𝒚. Here, we use the combined vector of LV and RV pressure and volume outputs as our 𝒚. We approximate the derivative using a centered-difference approach

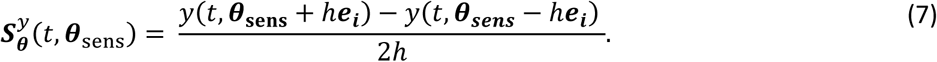

where 𝒆_𝒊_ is a unit vector in the *i*th direction. Given the various magnitudes of the parameters, we conduct sensitivity analysis on a log-scale, with ℎ = 0.01. Larger values of 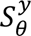 indicate greater sensitivity of the model to that parameter. To cover the parameter space, we performed the local sensitivity analysis on each of the Nx and Hx animals and their nominal parameters, corresponding to different values of 𝜽_𝐬𝐞𝐧𝐬_.

We normalized the sensitivities (i.e., 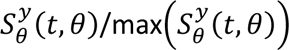) for each animal such that they varied from 0 to 1 (**Figure 2**), and the parameters with a strong effect (normalized sensitivity > 0.1) on the combined pressure and volume output (see **Equation 7**) were considered for additional analyses (Colunga et al., 2023). We used the sensitivity vectors to construct an approximate Fisher information matrix, 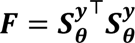, and ensured that for the selected parameter subset the condition number 𝑭 is less than 1e8 (Colebank et al., 2024).

**Figure 2.**
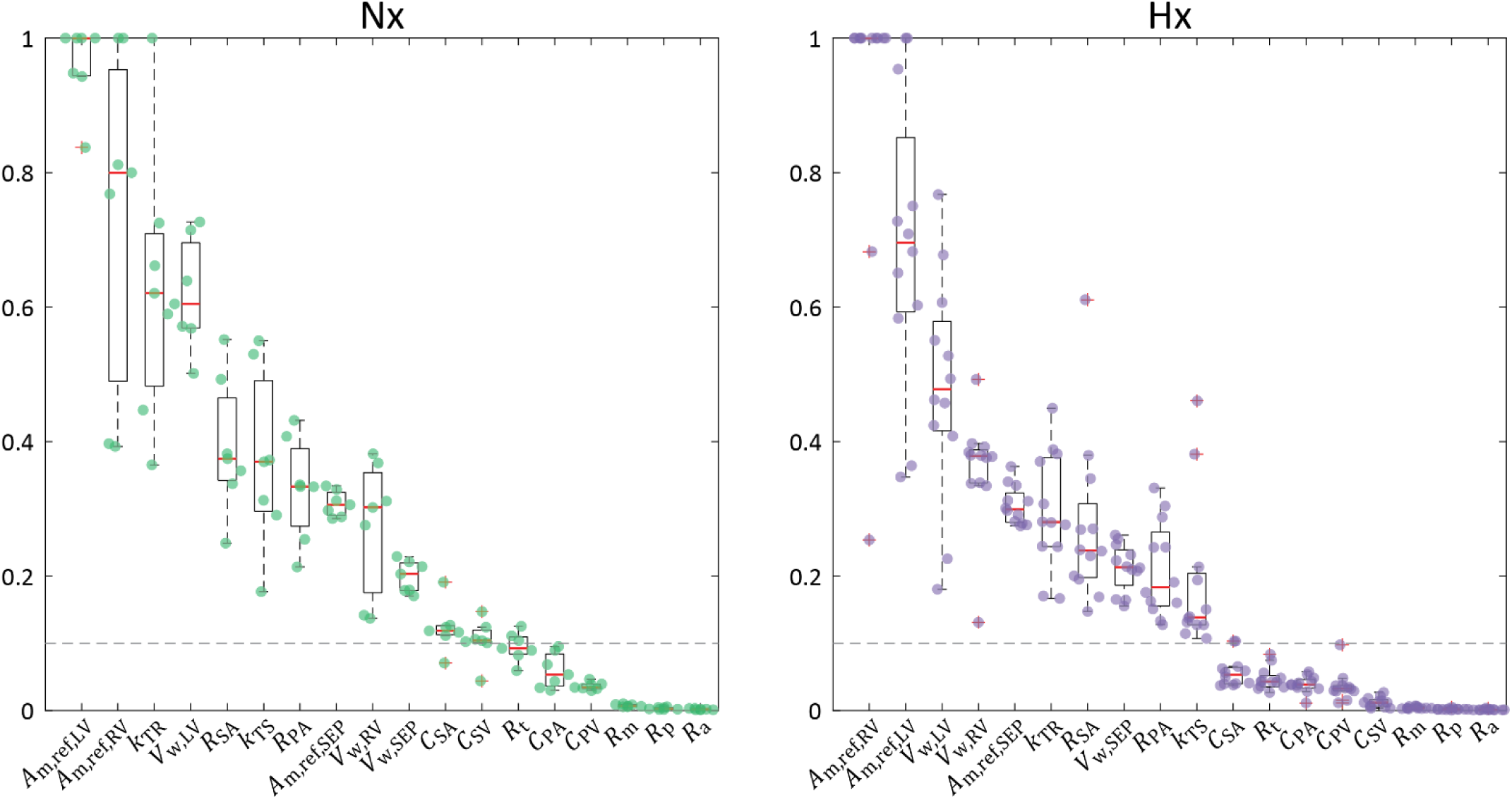
Sensitivity Analysis. Figure panels correspond to Nx (n = 7) and Hx (n = 12) experimental groups. The sensitivity of each parameter is normalized to each animal’s maximum sensitivity i.e. parameter sensitivity varies from 0 to 1. For each animal, the corresponding normalized sensitivity is indicated by a point. The box-and-whisker plots provide the interquartile range of the normalized sensitivy for each parameter in each experimental group (mean shown in red). Parameters are ordered as most to least influential based on the mean sensitivity ranking across all animals in the same experimental group. A threshold of 0.1 is prescribed to delinate between influential and non-influential parameters. The top 10 parameters for both Nx and Hx are above the threshold and are the same. Two additional parameters, *C*_SA_ and *C*_SV_, in Nx are also above the threshold.

### 2.6 Parameter Estimation and Model Calibration

From the sensitivity analysis, parameters with a normalized effect < 0.1 were fixed as non-influential. While all compliance parameters are below this threshold in Hx, *C*_SA_ and *C*_SV_ are slightly above the 0.1 threshold in Nx. Other than *C*_SA_ and *C*_SV_, all other parameters over the threshold were the same between Nx and Hx despite different orderings. We were interested in comparing parameter subsets between Nx and Hx and hence opted to use the same set of model parameters for each group and omitted *C*_SA_ and *C*_SV_ from the Nx group since they were only just over the threshold. Thus, our final parameter set for calibration for both Nx and Hx was

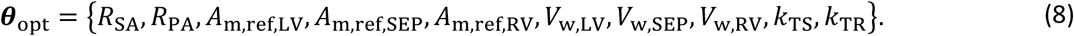

To calibrate the model, we minimized the cost function, 𝐽(𝜽_opt_) = 𝒓(𝜽_opt_) 𝒓(𝜽_opt_), where 𝒓 is the residual or difference between model simulations and measured data. We conducted our optimization on the log-scaled parameters to ensure similar magnitudes (Colunga et al., 2023). The components of 𝒓 include time dependent and scalar differences in the model and data. The four dynamic residuals for the volume and pressure waveform data of both ventricles, 𝒓_d_ = {*r*_V,LV_, *r*_V,RV_, *r*_P,LV_, *r*_P,RV_}, are given by

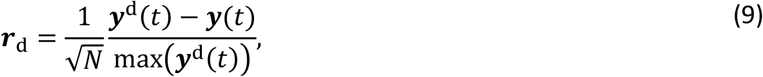

where 𝒚(𝑡) = {𝑽_LV_(𝑡), 𝑽_RV_(𝑡), 𝑷_LV_(𝑡), 𝑷_RV_(𝑡)} represents the model outputs and 𝒚^d^(𝑡) is the corresponding data. The model is resolved at timesteps that align with the waveform data, and the dynamic residuals are scaled based on the number of timesteps, 𝑁 = 50, in the data. The eight static residuals for the end-diastolic and end-systolic pressure volume for both ventricles are defined as

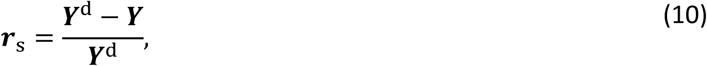

where the vector ***Y*** = {*EDV*_LV_, *EDV*_RV_, *ESV*_LV_, *ESV*_RV_, *EDP*_LV_, *EDP*_RV_, *ESP*_LV_, *ESP*_RV_} is the simulated pressure and volume, including end-diastolic and end-systolic volumes and pressures (EDV, ESV, EDP, and ESP), and ***Y***^d^ is the corresponding data. Therefore, the residual vector becomes 𝒓 = {𝒓_d_, 𝒓_s_}. We used gradient-based, nonlinear least-squares global optimization with 20 randomized starting parameter vectors selected between ± 50% of the log-nominal values to minimize 𝐽(𝜽_opt_) (Colunga et al., 2023). We chose 20 initial starts to provide a robust optimal fit to the data. Of the 20 optimized parameter sets we chose the set with the smallest cost, 𝐽, to use for continued analyses. The upper and lower for the parameters are provided in **Table 4**. The overall simulation protocol is summarized in **Figure 3**.

**Figure 3.**
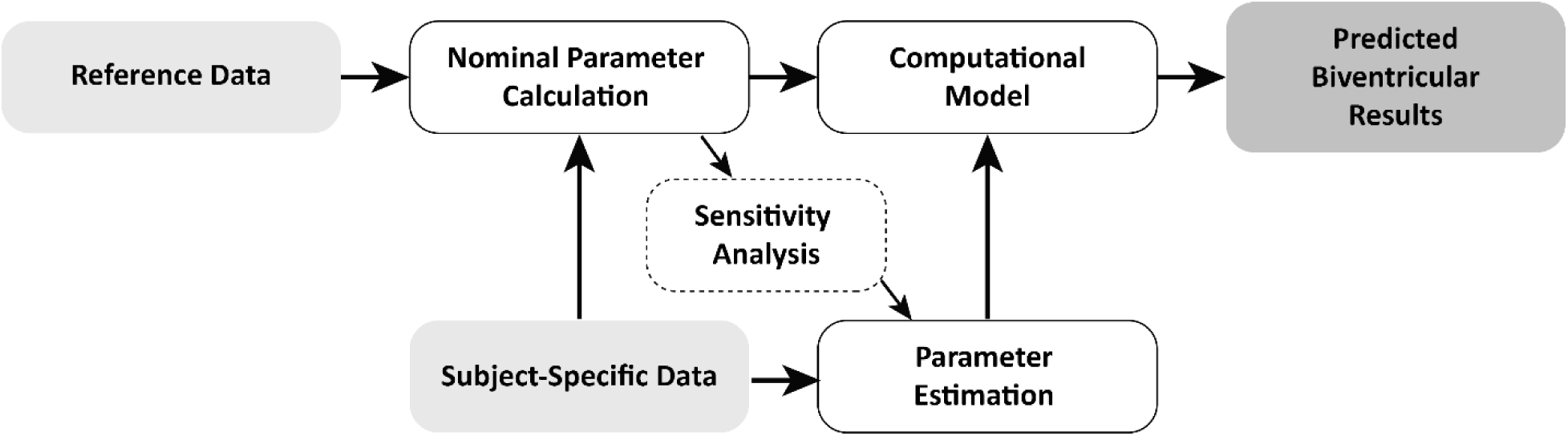
Simulation Protocol.

**Table 4.** Parameter Bounds for Model Calibration.

| Parameters ($\boldsymbol{\theta}_{\text{opt}}$) | Upper and Lower Bounds |
| --- | --- |
| $A_{m,ref,RV}, A_{m,ref,LV}, A_{m,ref,SEP}, V_{w,RV}, V_{w,LV}, V_{w,SEP}$ | $0.8 \cdot \log(\theta) < \log(\theta) < 1.2 \cdot \log(\theta)$ |
| $R_{SA}, R_{PA}$ | $0.1 \cdot \log(\theta) < \log(\theta) < 10 \cdot \log(\theta)$ |
| $k_{TS}$ | $0.001 < k_{TS} < 0.1$ |
| $k_{TR}$ | $0.3 < k_{TR} < 0.5$ |

$A_{m,ref,RV}, A_{m,ref,LV}, A_{m,ref,SEP}$ - midwall reference area for the right ventricle (RV), left ventricle (LV), and septum (SEP), respectively; $V_{w,RV}, V_{w,LV}, V_{w,SEP}$ - wall volume for the RV, LV, and SEP; $R_{SA}$ - systemic arterial resistance; $R_{PA}$ - pulmonary arterial resistance ; $k_{TS}, k_{TR}$ - proportion of cardiac cycle during systole and relaxation, respectively.

### 2.7 Computed Metrics for Comparison and Statistical Methods

We compared computed ESV, EDV, ESP, and EDP to the measured data. In addition, we quantified myofiber power intensity and relative sarcomere shortening. To assess septal dynamics, we computed the TriSeg septal curvature (Kim et al., 2023; Lumens et al., 2009).

Model-predicted end-systolic and end-diastolic points are the minimum and maximum volumes and the maximum and minimum pressures at those volumes, respectively. Relative sarcomere shortening is computed as the change in sarcomere length throughout the cardiac cycle relative to the sarcomere length at the start of the cardiac cycle, (𝑳_s,*i*_(𝑡) − 𝑳_s,*i*_(0))/𝑳_s,*i*_(0).

We approximated myofiber power intensity (power per unit area) by first computing myofiber work intensity (work per unit area) throughout the cardiac cycle with discrete approximation of the integration of active myofiber force per unit area, *σ*_XB,*i*_, with respect to sarcomere length, 𝑳_s,*i*_. Similarly, we approximated the derivative of the cumulative work intensity throughout the cycle with respect to time. We use the max absolute power intensity throughout the cardiac cycle as our comparison point across subjects.

We quantified septal “bounce” as the maximum drop in septal curvature during systole from the start of the cardiac cycle, Δ*C*_m,SEP_(cm^−1^). The average curvature during systole, 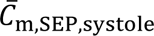, was computed for comparison of septal flattening, or lack thereof, between Nx and Hx groups. Stroke work, SW, is calculated as the area of the pressure-volume loop.

We used a student’s t-test to assess differences between 1) the computed model outputs and measured data, 2) the calibrated Nx and Hx parameters and model outputs and 3) the calibrated parameters for the Hx male and female animals (**Figure S1**). Additionally, we assessed monotonic relationships in the parameters using Spearman’s correlation coefficient (**Table 5**) and linear relationships between various modeling outputs using Pearson’s correlation coefficient (**Table 7**) and visualized the data via scatter plots (**Figure 7**).

**Table 5.** Comparison of Model Outputs to Corresponding Data.

| Metric | Units | Nx (n = 7) | | $R^2$ | Hx (n = 12) | | $R^2$ |
| --- | --- | --- | --- | --- | --- | --- | --- |
|  |  | Data | Model |  | Data | Model |  |
| LV EDV | $\mu\text{L}$ | **44**<br>$\pm 7$ | **44**<br>$\pm 8$ | **0.99** | **47**<br>$\pm 11$ | **46**<br>$\pm 10$ | **0.98** |
| LV ESV | $\mu\text{L}$ | **16**<br>$\pm 9$ | **16**<br>$\pm 10$ | **0.96** | **18**<br>$\pm 8$ | **18**<br>$\pm 9$ | **0.99** |
| RV EDV | $\mu\text{L}$ | **52**<br>$\pm 13$ | **51**<br>$\pm 14$ | **0.99** | **52**<br>$\pm 14$ | **51**<br>$\pm 13$ | **0.97** |
| RV ESV | $\mu\text{L}$ | **24**<br>$\pm 11$ | **24**<br>$\pm 12$ | **0.95** | **23**<br>$\pm 9$ | **23**<br>$\pm 10$ | **0.99** |
| LV EDP | mmHg | **4.4**<br>$\pm 2.0$ | **4.6**<br>$\pm 1.8$ | **0.98** | **5.1**<br>$\pm 1.4$ | **4.9**<br>$\pm 1.5$ | **0.97** |
| LV ESP | mmHg | **65**<br>$\pm 9$ | **65**<br>$\pm 10$ | **0.98** | **72**<br>$\pm 11$ | **68**<br>$\pm 11$ | **0.95** |
| RV EDP | mmHg | **2.9**<br>$\pm 1.5$ | **1.4**<br>$\pm 0.079$ | **0.56** | **3.8**<br>$\pm 1.5$ | **1.4**<br>$\pm 0.099$ | **0.14** |
| RV ESP | mmHg | **26**<br>$\pm 3$ | **26**<br>$\pm 3.2$ | **0.97** | **38**<br>$\pm 6$ | **34**<br>$\pm 8$ | **0.99** |

$R^2$ - Spearman's correlation coefficient.

We performed a linear discriminant analysis (LDA) to assess separability of the two conditions using three sets of features (**Figure 8**). The first set uses rat data alone, namely volume and pressure data described in **Table 1**. The second set uses the calibrated parameter values as described in **Figure 5**. The third and final set integrated both the preclinical rat data and the model-derived features.

## 3. Results

### 3.1 Model Calibrations

The calibrated pressure-volume loop simulations demonstrated good agreement with the measured data as shown in **Figure 4**. In general, model simulations were within the bounds of variability seen in the 20 or more heartbeats of data. The model had difficulty matching RV EDP (**Table 5**), with an error between 2-4 mmHg.

**Figure 4.**
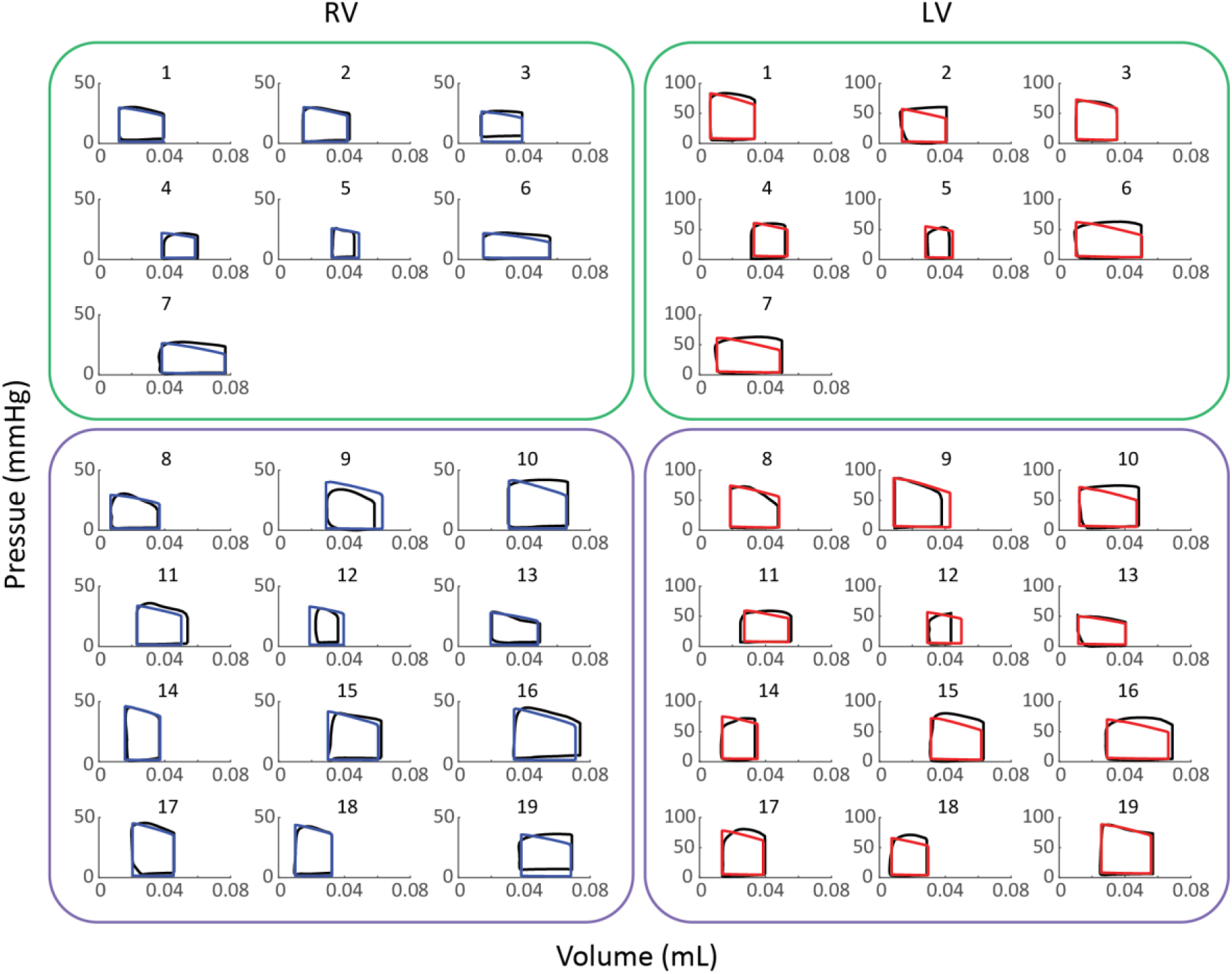
Calibrated Pressure Volume Loops. Calibrated Nx (green, subjects 1-7) and Hx (purple, subjects 8-19) subject-specific pressure volume loop simulations for the left ventricle (LV, red) and the right ventricle (RV, blue). The corresponding average of the data from 20+ cardiac cycles to which the model was calibrated is shown in black. The simulated pressure volume loops show good agreement with the measured data. The calibrated pressure and volume signals are shown in the Supplumenary Material.

Comparisons between the 10 model parameters (𝜽_opt_) calibrated to LV and RV pressure and volume data for Nx vs. Hx animals are shown in **Figure 5**. There are multiple significant differences in the calibrated parameters; the RV reference area, 𝐴_m,ref,RV_, and the systolic timing parameter, *k*_TS_, are significantly increased in Hx, and the septal wall volume, *V*_w,SEP_, is significantly decreased in Hx compared to Nx. Although the difference is not significant, the pulmonary arterial resistance, *R*_PA_, is notably increased in Hx (group mean for Hx: 24.3, for Nx: 16.7).

**Figure 5.**
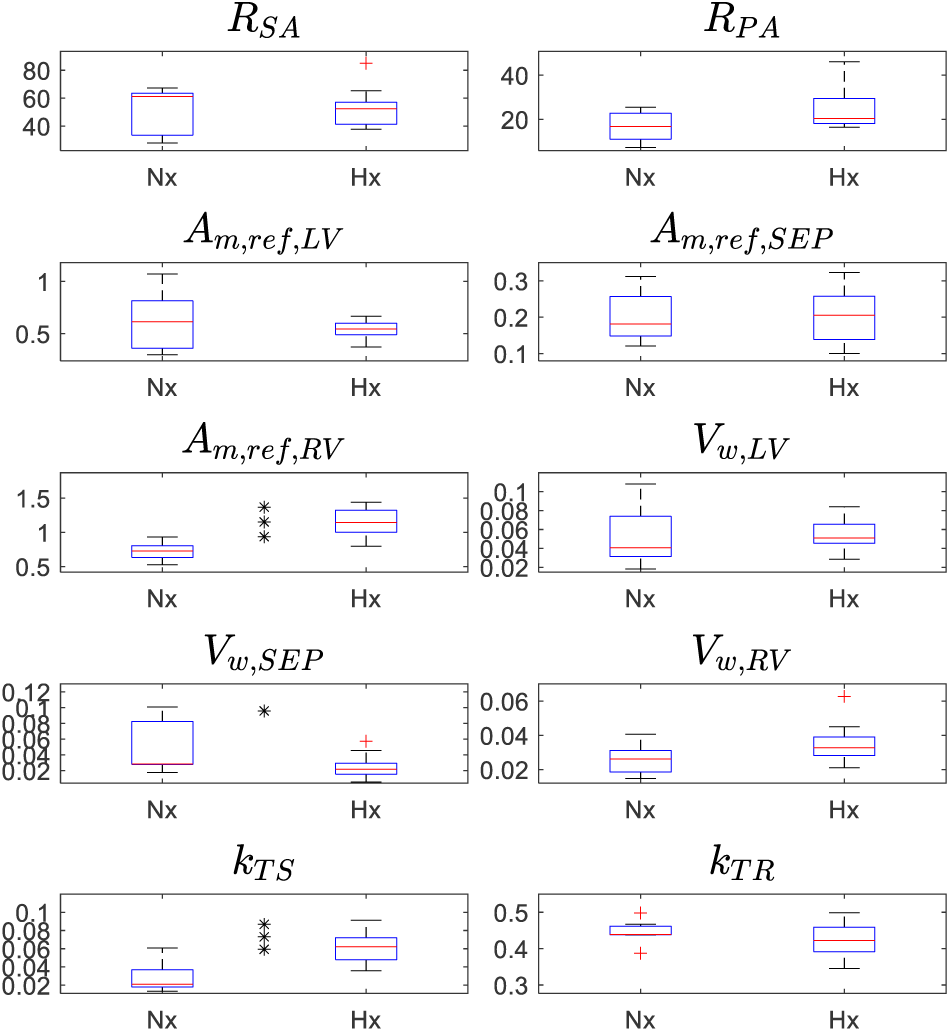
Calibrated Model Parameters (**𝜽_opt_**) for Nx and Hx. Comparison of calibrated model parameter distributions for Nx and Hx, where 𝜽_opt_ = {*R*_SA_, *R*_PA_, 𝐴_m,ref,LV_, 𝐴_m,ref,SEP_, 𝐴_m,ref,RV_, *V*_w,LV_, *V*_w,SEP_, *V*_w,RV_, *k*_TS_, *k*_TR_}. * Indicates a significant difference from an unpaired two-sample student’s t-test (p<0.05); ** (p<0.01); *** (p<0.001).

We compare the changes in calibrated parameters for each condition, separated by sex (**Table S14**). There is an insufficient number of female, Nx animals (Nx-F: n = 2) for statistical analysis, but *R*_SA_ and *R*_PA_ are notably increased in female Hx (66% and 135% increase in the means, respectively). Between the Nx and Hx males there are significant increases to 𝐴_m,ref,RV_ and *k*_TS_, which aligns with sex-pooled results. Interestingly, there are no significant differences in *V*_w,SEP_ for the males (Nx-M: 0.0299, Hx-M: 0.0297); in this case the finding in the sex-pooled group can be attributed to the female subjects (Nx-F: 0.0973, Hx-F: 0.0173). When we consider sex differences in the parameters for each condition, we observe that *R*_SA_ and *R*_PA_ are significantly greater for the Nx males (116% and 140% increase in the means, respectively) than the females, with no significant differences between the Hx males and females. Additionally, *R*_PA_ and *V*_w,SEP_ were markedly greater for the Hx males (41% and 72% increase in the means, respectively) compared to the females.

The computational model takes on average 1-2 seconds on a laptop to provide simulation results. This time can increase to up to a minute of computation time if the parameterization leads to stiff dynamics in the systems, issues with solving with differential algebraic equations, or reaching steady state. Optimizing all 20 initial parameter estimates takes between one to four hours when run in parallel on a six-core machine. We use a trust-region-reflection algorithm in the lsqnonlin.m function from MATLAB with function evaluations limited to 2000 but otherwise use the default settings. The tolerance for parameter changes, function evaluation, and the step size is set to 1e-6. Finally, the objective function values are calculated using the residual functions in equations (9-10). The objective function values for the optimal solutions are on the order of 10^−1^.

In effort to circumvent identifiability issues and local minima in our objective function, we combine our local sensitivity analysis with a multistart optimization as described in **Section 2.6**. The 20 optimized parameters cluster close to the optimal parameter set, but the parameters are numerically unique, and small differences in parameter values lead to differences in pressure and volume output signals. This outcome suggests that the model outputs are unique for each parameter combination and is consistent with Colebank and Chesler (2022), where a more in-depth identifiability analysis was done on a similar model using the TriSeg infrastructure. We credit the identifiability of the model parameters in this study to the high-fidelity pressure-volume data.

### 3.2 Model Predictions

As expected, mean pulmonary arterial pressure (mPAP) was elevated in Hx (**Table 6**). There was no difference in RV myofiber power intensity for the Hx group compared to the Nx group (**Table 6**) despite a prescribed increase in myofiber stiffness attributed to Hx (**Table 3**). However, the absolute RV myofiber length averaged over the cardiac cycle, 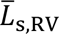, was lower in Hx than in Nx (**Table 6, Figure S2**), which suggests less engagement through the Frank-Starling Mechanism. The increased 𝐴_m,ref,RV_ in Hx (Figure 5) corresponds with the reduction in 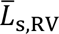.

**Table 6.**
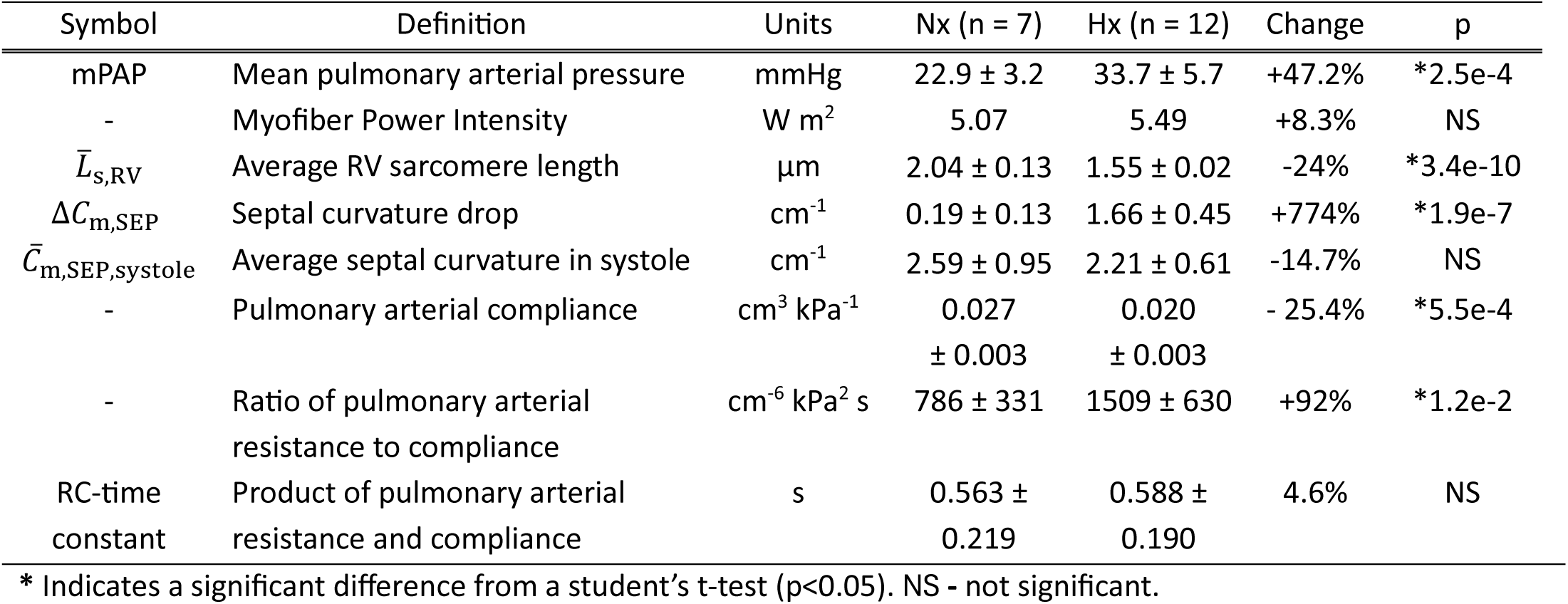
Nx vs Hx Metrics.

Model simulations revealed abnormal septal behavior during systole in Hx characterized by a rapid decrease (flattening towards the LV) and then subsequent rapid increase to baseline septal curvature near the start of ejection, or septal bounce (Figure 6). The septal drop, Δ*C*_m,SEP_, in Hx was nearly 8 times larger than in Nx (**Table 6**). The mean curvature over systole, 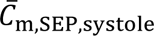, was lower in Hx than Nx, although this difference was not significant (**Table 6**). The nature of the bounce includes an almost mirrored return to normal baseline septal curvature after the initial rapid curvature drop. Thus, there were minimal differences in 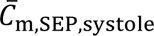 between Nx and Hx (**Table 6**).

**Figure 6.**
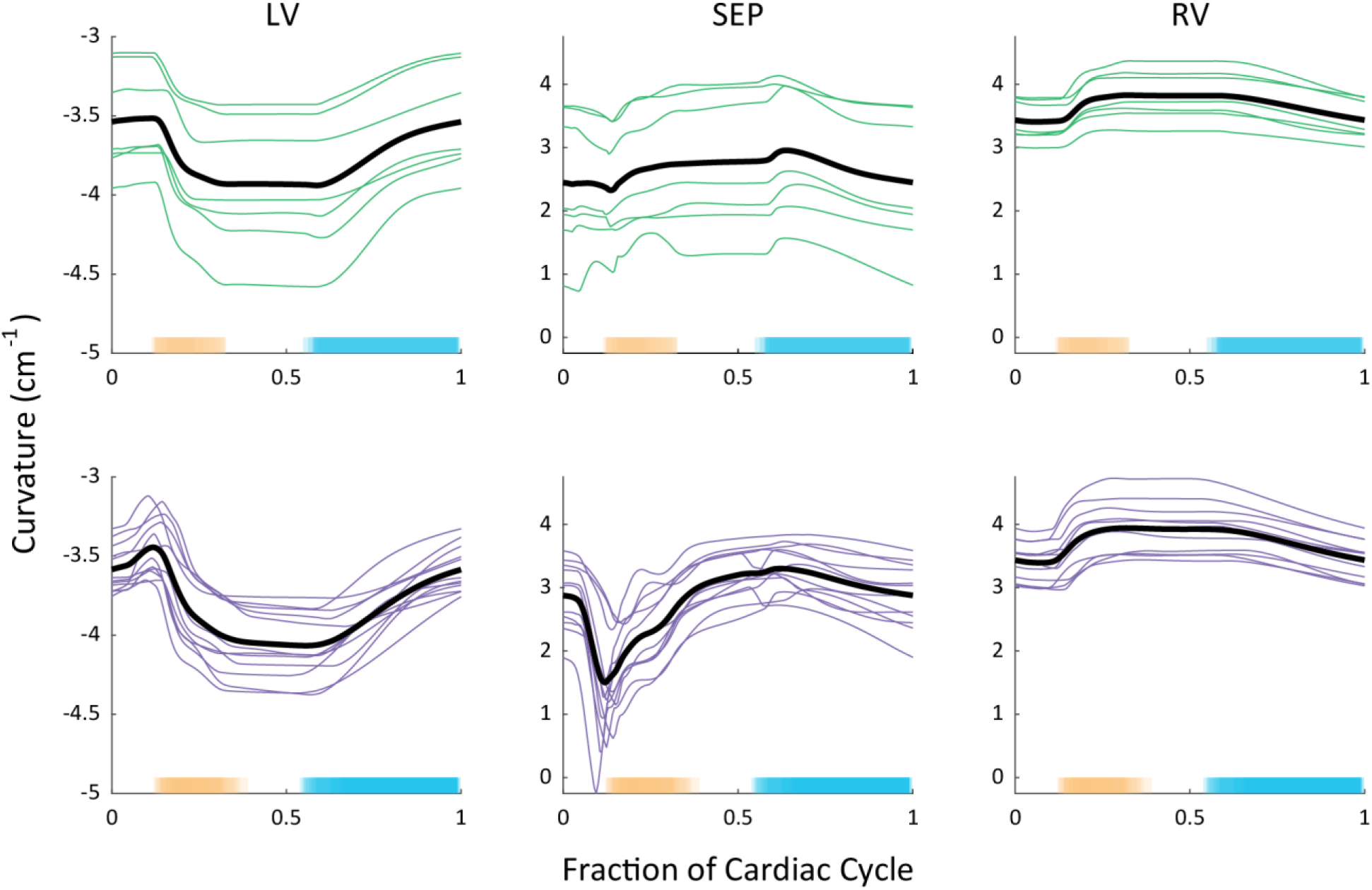
Curvature Predictions. Septal curvature from the start of the cardiac cycle for individual Nx and Hx animals. The x-axis is time-normalized to one cardiac cycle and the individual animal data is averaged (black) for ease of comparison. A lower septal curvature indicates flattening of the septum towards the LV. LV - left ventricle; RV - right ventricle.

We calculated the ratio of pulmonary arterial resistance to compliance as a metric of afterload and their product, the RC-time constant. The resistance-compliance ratio captures the effect of both altered resistance and compliance as contributors to afterload. Their ratio nearly doubled in Hx, which aligns with expected increases to right ventricular afterload. The RC-time constant is maintained across the conditions, consistent with clinical data in the pulmonary circulation (Tedford et al., 2012).

We performed a Pearson’s Test for linear relationships between multiple model outputs and parameters (**Table 7**) and visualized these relationships in Figure 7. The clinical measure, mPAP, has a significant and moderate positive relationship with the model-predicted septal curvature drop, Δ*C*_m,SEP_. In turn, the model-predicted Δ*C*_m,SEP_ has a significant and moderate positive relationship with metrics associated with RV function, including RV SW and 𝐴_m,ref,RV_.

**Figure 7.**
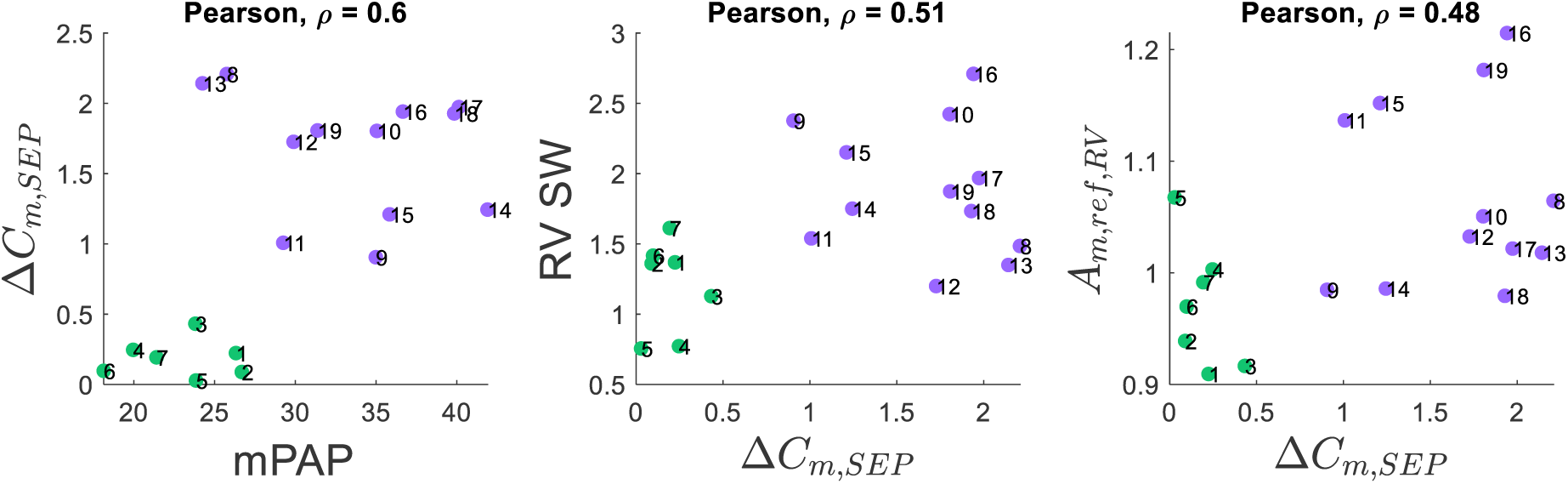
Relationships Between Model Outputs.

**Table 7.** Pearson’s Tests for Linear Relationships.

| Input 1 | Input 2 | $\rho$ | p |
| --- | --- | --- | --- |
| $\Delta C_{m,SEP}$ | RV SW | 0.51 | *1.2e-3 |
| $\Delta C_{m,SEP}$ | mPAP | 0.60 | *5.9e-5 |
| $\Delta C_{m,SEP}$ | $A_{m,ref,RV}$ | 0.48 | *2.8e-2 |

Correlation ( $\rho$ ) coefficient and significance (p) for linear relationships. \*Indicates a significant linear relationship (p<0.05).

Visualization of the LDA results, Figure 8, demonstrate increased separability with the inclusion of model-derived features compared to rat data alone. The analysis applied to the data alone results in partial overlap between the Nx and Hx groups and therefore limited separability. However, the LDA performed on the calibrated model parameters (Figure 5) alone are clearly separated, and the separation increases when the rat data and model parameters are combined. This suggests that model-derived parameters, including those representing unmeasurable physiological biomarkers, can provide additional discriminatory information.

**Figure 8.**
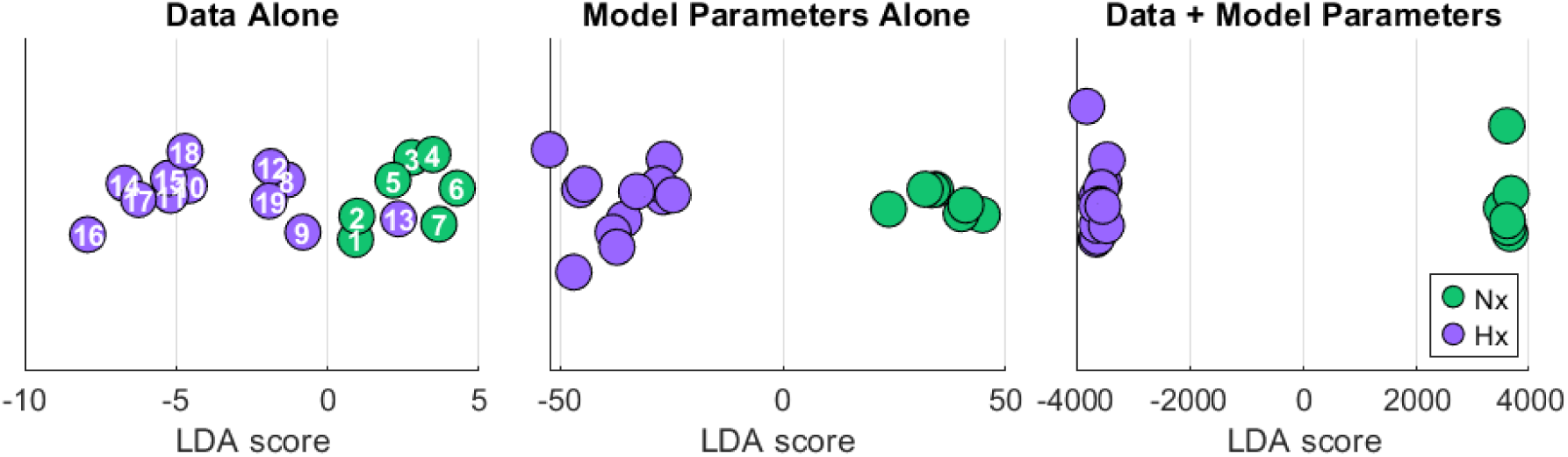
Linear Discriminant Analysis. LDA projections demonstrating separability of Nx and Hx conditions using rat data, calibrated model parameters, and combined data and model parameters. The combined feature set demonstrates the greatest amount of separability between the conditions.

## 4. Discussion

In this study, we developed subject-specific, multiscale models of the cardiopulmonary system in rodents that demonstrated an excellent fit to the experimental data and captured cardiopulmonary abnormalities that are consistent with the clinical literature on preterm born subjects. Our results highlight the multiscale nature of the cardiopulmonary consequences of postnatal Hx with implications for preterm born individuals and provide a framework for further exploration of cardiopulmonary disease manifestation into adulthood.

### 4.1 Simulation and Sensitivity Analysis

Versions of the multiscale cardiac model (Marzban et al., 2020) and TriSeg model (Kim et al., 2023) have demonstrated the ability to capture physiological hemodynamics in adult rats and humans, but the model has not been previously calibrated to pediatric and Hx cardiopulmonary data; therefore, we applied established approaches for model calibration that are scalable and promote identifiability. Colebank and Chesler (2022) demonstrated that in the context of biventricular, multiscale models for the study of pulmonary hypertension and RV function, right heart catheterization alone still yields identifiability issues and that at least some LV data is important to obtain. In this study, we had access to both RV and LV catheterization data; however with over 50 parameters in the full multiscale model and limited cellular-scale data, all parameters cannot be inferred simultaneously (Colebank et al., 2024).

To overcome this, we fixed parameters at the cellular-scale based on prior Nx rodent models (Marzban et al., 2020) and literature data in preterm birth and neonates (Patel et al., 2017; Prakash et al., 1999). We then used local sensitivity analysis to identify the most influential parameters for model calibration. Our findings in Figure 2 show similar parameter effects in both Nx and Hx animals. However, the sensitivity of pressure-volume simulations to RV reference area increased in Hx, with nearly all Hx animals showing 𝐴_m,ref,RV_ as the most important parameter. Conversely, Hx sensitivity to LV reference area was smaller than Nx, again suggesting a shift in dominance between the two ventricles. Similar conclusions can be made for the wall volumes, *V*_w,RV_ and *V*_w,LV_. These shape parameters have been identified as highly influential in other modeling studies (Colebank & Chesler, 2022; Jones & Oomen, 2025). In general, valve resistance and vascular compliance parameters were the least influential on pressure and volume outputs.

Our approach parallels methods employed by computational preterm studies combining physics-based mathematical modeling with subject data. For example, May et al. (2023) combined clinically measured cardiovascular anatomical and functional data from infants with a closed-loop computational model of the neonatal circulation. Subject data was partitioned for direct input into the model, parameter estimation, and verification of model predictions. For our study, more data is required for verification of model predictions as discussed in the limitations (**Section 4.7**).

### 4.2 Pressure-volume Loop Calibration

End-diastolic and end-systolic pressures and volumes have greater relevance than the intermediate points in the cardiac cycle, thus we put greater relative weight on the static end-systolic and end-diastolic measures in the calibration cost function. With this approach we were able to reach a global minimum across multiple randomized starting points, and the calibrated model simulations predict subject specific pressure-volume dynamics within the bounds of variability in data. A similar weighting approach was used by Colunga et al. (2023), which considered dynamic pressure data in the right heart and pulmonary artery along with static pressure data. The authors concluded similarly that static data corresponding to systolic and diastolic function were informative for parameter estimation, but that timeseries data provided additional insight. Our model consistently under-predicted RV EDP by 2-4 mmHg (**Table 5**).

However, overall, the group averages of the calibrated model outputs were consistent with the corresponding group averages of the measured data. Note that we limited the number of model parameters to prioritize identifiability and interpretability of the parameters rather than include more non-identifiable parameters that provide a better fit at the expense of this interpretability. The ability of the model to match organ-level dynamics for individual subjects without detailed information on microscale mechanisms is a strength of our study.

Subject-specific modeling allowed us to compare the relationships between multiple model-predicted outputs and parameters, which can help determine relationships between multiple clinically relevant metrics (Figure 7, **Table 7**). Additionally, findings from our LDA analysis (Figure 8) suggest that incorporating mechanistic modeling can provide further insights into disease mechanisms that are not captured through preclinical measures alone. With more data, unsupervised machine learning may help stratify subjects into treatment cohorts based on non-invasive measures and model outputs (Gu et al., 2025; Jones et al., 2021), however the present dataset is too small for robustness in these analysis techniques. Given this limitation in experimental data, the mechanistic model could be used to generate synthetic data for training of these machine learning models as in Jones and Oomen (2025), although that is beyond the scope of this work.

### 4.3 Mechanisms of RV Hypercontractility

Our model investigations indicate multiple mechanisms by which myofiber contractile function may be altered from Nx to Hx. At least six parameters in the crossbridge model modulate Ca^2+^ sensitivity and maximum Ca^2+^ activated force within physiologically relevant ranges (**Table 2**). As detailed in our analysis in **Table 3**, the combination of decreasing the rate of Ca^2+^ unbinding from Troponin C via *k*_off_ and increasing *k*_stiff,1_ and *k*_stiff,2_ yields the expected changes in Ca^2+^ sensitivity and maximum Ca^2+^ activated force. Equivalent increases to Ca^2+^ binding to Troponin C rate *k*_on_ and decreases to *k*_off_ yield the same steady-state increase in Ca^2+^ sensitivity as assessed by the force-pCa curve (Chung et al., 2016; Dowrick et al., 2023; Kreutziger et al., 2011; Saad et al., 2023), which is also reflected in the model. However, there are distinct differences in the dynamic force development and relaxation of the myofiber when *k*_on_ versus *k*_off_ is altered (Chung et al., 2016). Namely, an increase in Ca^2+^ sensitivity due to decreased *k*_off_ yields slower relaxation kinetics than increased *k*_on_. To confirm the mechanism of increased RV myofiber power with Hx and/or preterm birth, dynamic Ca^2+^ sensitivity experiments (Chung et al., 2016) could be used to measure *k*_on_ and *k*_off_. Thus, our multiscale model provides a viable approach for testing multiple physiological hypotheses that may drive changes in the RV during postnatal Hx exposure. This innovative integration of data across multiple studies enables a robust testing of established hypotheses that is typically infeasible using experimental data alone.

RV myofiber power intensity is not significantly changed in Hx compared to Nx (**Table 6**) despite increased RV SW and directly imposed RV myofiber contractility changes (as in **Table 3**) to mimic experimental observation in Hx. However, there are numerous mechanisms that contribute to contractile function (Sharifi Kia et al., 2021), and this mismatch suggests additional mechanisms contributing to unchanged myofiber power intensity yet increased SW. For example, the time fraction for the rise of calcium development, *k*_TS_, is significantly increased in Hx, which suggests an increased time for contraction contributes to overcoming the increased afterload in Hx. While RV myofibers are more contractile in Hx, the absolute RV sarcomere length was significantly reduced in Hx when compared to Nx, which corresponds to less engagement through the Frank-Starling mechanisms. Direct measurement of sarcomere length in whole organs is possible (Botcherby et al., 2013) but was not performed here to validate these predictions. Thus, direct translation of model sarcomere length to in vivo measures should be made with caution. In the model, sarcomere length is dependent on ventricular wall area and therefore represents the mechanical changes experienced by the ventricular wall. Together, these observations suggest a less efficient RV, which warrants further exploration.

### 4.4 Abnormal Septal Dynamics as a Marker of Altered RV Contractile Function and Pulmonary Hypertension

In our study, average septal curvature during systole (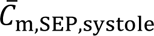) did not significantly decrease in Hx despite significantly increased mPAP (**Table 6**). We attribute this result, at least in part, to the nature of septal bounce phenomenon wherein the curvature rapidly decreases, but then just as rapidly increases to return to baseline curvature. Clinically, assessment of septal flattening can be highly variable (Mourani et al., 2015), however septal flattening has been observed in echocardiographic studies of nearly 300 preterm infants at P7 with pulmonary hypertension (Mourani et al., 2015) and over 200 infants at risk for pulmonary hypertension with systolic septal flattening as quantified by an LV end-systolic eccentricity index above 1.3 (Abraham & Weismann, 2016).

While average septal curvature over systole (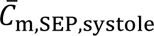) was a poor metric for differentiation between Hx and Nx in our simulations, our results showed significant correlations between septal bounce during systole and RV dilation as well as septal bounce during systole and RV SW (**Table 7**), which corresponds with observed correlations between flattening and RV dilation (Mourani et al., 2015) and flattening and RV systolic dysfunction (Abraham & Weismann, 2016). A previous study found that rapid leftward septal motion (equivalent to Δ*C*_m,SEP_) during early LV diastole is a marker of worsening RV function in chronic pulmonary arterial hypertension resulting from interventricular relaxation desynchrony and altered RV contraction (Palau-Caballero et al., 2017). While our subject population is distinct from that of Palau-Caballero et al., their findings demonstrate the utility of septal bounce as a metric for altered contractile function. Therefore, we suggest that systolic septal bounce may be a more sensitive noninvasive marker of altered RV contractile function and acute pulmonary arterial hypertension than absolute flattening in pediatric subjects. That said, the complexities in the multiscale model make it difficult to isolate the exact reason for changes in curvature and warrant further investigation. Future experiments that report cardiac wall shortening and dynamics prior to ex-vivo tissue analysis may reveal the multiscale links between RV function, septal motion, and cellular or subcellular processes.

### 4.5 Sex Differences

In the preclinical dataset, the female Hx rats had increased RV ESV compared to the males (mean of 29 µL vs 19 µL), which was reflected in the model, but otherwise there were no significant differences between the males and females in the data or in calibrated model parameters (**Table S1, S14**). That said, pulmonary arterial resistance, *R*_𝑝𝑎_, (Hx-F: 19.6, Hx-M: 27.6) is dramatically higher in the males. Interestingly, the greater pulmonary arterial resistance in males occurs despite comparable RV ESP (Hx-F: 36, Hx-M: 39). We also note a smaller stroke volume in males versus females, which may explain part of this difference given that *R* ∝ Δ*P*/*SV*. This greater resistance suggests a worse cardiopulmonary status and corresponds with clinical findings that preterm neonates are more often male, and males exhibit worse outcomes and greater death rates than females (Goss et al., 2020; Ingemarsson, 2003).

In the rat Hx model of preterm birth, sex differences at 1 year have been identified (Cantu et al., 2023); at P21, the age considered in this study, the observed sex differences thus far are limited (Kumari et al., 2019; Rahmani et al., 2024). Therefore, further mechanisms of sex differences may be identified by modeling rat Hx data at 1 year (P365) or by including sex hormones in a growth and remodeling framework (Lakshmikanthan et al., 2025).

### 4.6 Implications for Human Preterm Birth Modeling

Modeling and simulation of the cardiovascular system is a crucial component of precision medicine, especially with the development of medical digital twins (Colebank et al., 2024). The primary goals of cardiovascular systems modeling are two-fold: 1) to augment clinical and experimental data for knowledge generation, as in this study, and 2) to be a clinical tool for diagnosis and treatment. It should be noted that modeling approaches typically focus on calibration and validation to adult patients, whereas this approach is relatively limited in its application in pediatrics. When developing tools for clinical use a key consideration is the data availability (Niederer, Lumens, et al., 2019).

In this study, dynamic pressure and volume data were available for model calibration, but in a clinical setting the available data is likely to be far more limited (e.g., systolic and diastolic metrics via echocardiography) and require a modification of our approach. The timepoint of the rats in this study corresponds to human early childhood, from whom it is particularly challenging to collect data. Versions of the mechanistic model have been parameterized for both adult humans, mice, and rats, and now, young rats; therefore, it is reasonable to expect that the model can be applied to infant humans in the future. Depending on the age of the subject, data collected in those born preterm can include systemic blood pressure, heart rate, oxygen saturation, major vessel diameters (via MRI), blood velocity profiles (via ultrasound), and cardiac geometry and morphology/tissue properties (such as fibrosis, via cardiac MR) (Barton et al., 2021; Philip A. Corrado et al., 2021; P. A. Corrado et al., 2021; François et al., 2022; Kumari et al., 2021; May et al., 2023). In young adults and adults, right heart catheterization may be justified (Goss et al., 2018), but in infants this invasive method is typically only used in cases of suspected severe pulmonary hypertension, congenital heart defects or when surgery is already required. Taking these factors into consideration, it is reasonable to consider that a typical clinical dataset would include cardiac volumes and geometry (area, thickness), static systemic blood pressure estimates (subsequently LV pressure), static cardiac pressure estimates, total blood volume estimates, and estimated time spent across cardiac stages. However, the more limited dataset may not be sufficient for robust model calibration (Colebank & Chesler, 2022), in which case the addition of statistical models may be useful or necessary to derive meaningful insights (Gu et al., 2025; Peirlinck et al., 2021). Echocardiography is often the first noninvasive step to determine disease severity and the possible need for catheterization (Mourani et al., 2008), thus one application of modeling could be to reduce the uncertainty in echocardiography-based predictions (Mourani et al., 2015) and aid in the identification of high-risk patients who would benefit from early intervention.

At a microscale level, major components of contractile function are maintained across species; however, humans express primarily beta-myosin heavy chain (MHC) isoforms with small amounts of alpha-MHC in the ventricles while the reverse is true in rats (Prodanovic et al., 2022). Therefore, implications derived from analyses of contractile function in early life Hx rats for preterm humans must be considered with caution (Niederer, Campbell, et al., 2019; Reiser et al., 2001). A key difference between the two MHC isoforms is in the transition rates between various crossbridge states. The 5-state crossbridge model presented in this study includes rate transitions that can be tuned to data, and future work could include adjusting crossbridge kinetics (Johnson et al., 2019; Prodanovic et al., 2022) to account for differences in alpha and beta isoforms, allowing for improved translation of experimental rodent data to human subjects (Niederer, Campbell, et al., 2019).

### 4.7 Limitations

The major limitations of this study include limited mitochondrial and crossbridge-scale data, lack of imaging data, and a small subject population. As a result, validation of many model parameters, especially at the cell and tissue level, must await additional data. Collection of subject-specific crossbridge data would allow for model calibration at the cellular-scale and greater insights into cellular mechanisms of increased myofiber power with Hx. This data would also narrow the possibilities of Hx crossbridge dynamics, as outlined in **Table 3**. Understanding the connection between energetics and mechanics is also important for parameterizing and calibrating the model and requires further study and data collection. Additionally, wall weights or volumes for each animal were not available and we used an approximation based on group averages. Access to subject-specific wall geometry data would improve our ability to further personalize the model. We did not have access to imaging data, such as echo or MRI, to confirm the abnormal septal motion in Hx that the model predicts. However, our findings correspond with clinical studies in preterm infants and infants with pulmonary hypertension and computational studies of pulmonary hypertension, providing support for our observations. Though our study includes pressure-volume data from both ventricles, we find that the end-diastolic pressure-volume relationship is relatively flat, hence our model results to do not show nonlinear EDPVR dynamics. Indeed, the data also exhibits a flat EDPVR, thus the model and data are aligned in this regard. Finally, the small subject population prevented dividing the dataset into calibration and validation cohorts, which is considered a best practice (Colebank et al., 2024) and limited our ability to interrogate sex differences (n=2 females in the Nx group).

## 5. Conclusions

By calibrating a multiscale cardiac mechanics and closed loop circulation model to data from an established rodent model of preterm birth, we highlight key cardiovascular consequences of postnatal Hx exposure during early life. We provide a framework that can take complex, multiscale and multiorgan models, and parameterize them such that disease mechanisms can be tested, and intrinsically subject-specific parameters are calibrated to available data. Moreover, our study is the first to consider a subject-specific, multiscale cardiopulmonary model for analyzing the effects of postnatal Hx in rodents. Our investigations generate testable hypotheses regarding mechanisms of a hypercontractile RV due to postnatal Hx and lay the foundation for future studies on the cardiopulmonary consequences of preterm birth in humans.

## 6. Supplementary Material

With this manuscript all model codes are provided, available at https://github.com/sallakim/Multiscale-Hx-P21-Model-2025.

## Acknowledgements

National Institutes of Health, National Heart Lung and Blood Institute R01 HL154624 (NCC, DAB) National Institutes of Health, National Heart Lung and Blood Institute R01 HL173346 (DAB)

MJC was supported in part through TL1 TR001415 through the National Center for Research Resources and the National Center for Advancing Translational Sciences, National Institutes of Health (NIH).

Original animal studies used for modeling were supported by the University of Wisconsin Clinical and Translational Science Award program, through National Center for Advancing Translational Sciences Grant UL1-TR-000427 (primary investigator M. Drezner; 4KL2-TR-000428-10, awarded to KNG), as well as a Joel Belt Pediatric Pulmonary Hypertension Research and Mentoring Grant through the Pulmonary Hypertension Association (Barst Award; PI KNG).

## Statements and Declarations

Competing Interests: No conflicts of interest to disclose.

